# Alpha oscillations link action to cognition: An oculomotor account of the brain’s dominant rhythm

**DOI:** 10.1101/2021.09.24.461634

**Authors:** Tzvetan Popov, Gregory A. Miller, Brigitte Rockstroh, Ole Jensen, Nicolas Langer

**Affiliations:** Department of Psychology, University of Zurich, Methods of Plasticity Research, Department of Psychology, University of Zurich, Zurich, Switzerland; University Research Priority Program (URPP) Dynamics of Healthy Aging, Zurich, Switzerland; Neuroscience Center Zurich (ZNZ), Zurich, Switzerland; Department of Psychology and Department of Psychiatry and Biobehavioral Sciences, UCLA, USA; Department of Psychology, University Konstanz, Konstanz, Germany; School of Psychology, University of Birmingham, Birmingham, UK

## Abstract

Power modulations in alpha oscillations (8-14Hz) have been associated with most human cognitive functions and psychopathological conditions studied. These reports are often inconsistent with the prevailing view of a specific relationship of alpha oscillations to attention and working memory (WM). We propose that conceptualizing the role of alpha oscillations in oculomotor control resolves this inconsistency. This proposition is based on a review of results across species (human N_pooled_=295, one non-human primate, honey bee N=5), experimental conditions (rest, attention, and working memory), and recording techniques (EEG, ECOG, eye-tracking, and MEG) that encourage the following relationships between alpha oscillations and eye-movement control: (i) saccade initiation prompts power decrease in brain circuits associated with saccadic control; (ii) the direction of a saccade is consistent with alpha lateralization, both during task and resting conditions; (iii) the phase of alpha activity informs saccade occurrence and biases miniature eye movements during fixation (e.g. fixational tremor); and (iv) oculomotor action differentiates WM load. A new theory on how alpha oscillations link oculomotor action to cognition is proposed. Generalizing across tasks and species: low oculomotor activity is associated with high alpha power and vice versa. Alpha oscillations regulate *how long* to look at a given target and *how fast* to saccade to a next. By ensuring steady gaze position, any potential input outside foveal vision is “suppressed”.

## Introduction

Alpha oscillations refer to the dominant rhythm of the human brain in the frequency spectrum, approximately 8-14 Hz. Since Berger first defined this dominant rhythm and assigned it to “relaxed waking”, the functional significance of the rhythm and its modulation during information processing and behavior control have been core issues prompting extensive research and controversial discussion. Here, we propose a crucial role of alpha oscillations in oculomotor control as a prerequisite of cognitive-behavioral processing. In the following, existing and then new empirical and experimental evidence in support of this oculomotor theory of alpha oscillations is discussed, followed by an elaboration of the theory.

Eye movements are produced continuously, typically expressed as saccades that occur at the rate of 3-4 Hz^1^, as well as micro-saccades with comparable spatiotemporal dynamics^2^. During fixation periods, eye movements do not actually stop. Instead, micro-saccades^2^ and eye movements at a rate of 40-90 Hz continue (described as physiological nystagmus or fixational tremor^3^). Their omnipresence during waking, including cognitive tasks, suggests a mechanism that mediates between the various types of eye movement such as macro- and micro-saccades as well as fixational tremor. A first indication for a basic relationship between the dominant alpha rhythm and oculomotor action is that this relationship requires calibration as it develops^4^. Neonates lack an alpha rhythm up to the 9^th^ month^5^. The peak frequency and power of the rhythm follow a developmental trajectory reflected in an increase in peak frequency and power with an increase in age^4^. Moreover, the fact that a dominant rhythm is seen in patients in a locked-in state with residual ability to communicate via eye movements but not in those without any means of communication supports the relationship between alpha oscillations and oculomotor functions^6, 7^. Further support for a relationship between the dominant alpha rhythm and oculomotor action is the more recent finding that the dominant rhythm allows the prediction of the covertly attended location in the subject’s environment^8–11^. That is, the spatial topography of alpha power at the scalp codes the position of a stimulus in the external visual field (henceforth we refer to this phenomenon as place topography). However, observations reporting that the locus of attention can be predicted solely on the basis of eye movements and independent of visual input ^12–15^ have led to caution about the mechanistic role of alpha power modulations in spatial attention. Is the external stimulus or covertly maintained spatial location the initiator of alpha power modulation? To address this question, consideration of the neural circuit controlling saccadic eye movements appears necessary.

Eye movements are initiated by activity originating in brainstem (BS) neurons of the abducens, trochlear, and oculomotor nuclei^16, 17^ (for overview see also^18, 19^). The firing pattern of the neurons in these nuclei determines the size of the saccade or the maintenance of object position on the retina during smooth pursuit^17, 20^ - higher firing rate coincides with larger saccade^21^. These BS nuclei are innervated by neurons residing in the superior colliculus (SC). The SC is anatomically organized in multiple layers and functionally subdivided in superficial and deep layers. The superficial layers are innervated directly by the retina and are subject to a locally wired center-surround antagonism, with these neurons responding robustly to stimuli occupying 0.3°-5.2° degrees of visual angle^17, 22^. Thus, stimuli from the external environment, such as a small bright spot stimulating the retina, are capable of triggering a prosaccade directly via the superficial SC layers.

Yet the environment can be explored, and saccades can be generated in the absence of external salient input. This is enabled by cortico-collicular connections that involve neurons in the deep layers of the SC. Crucially, the firing rate of deep-layer SC neurons does not inform saccade initiation. Instead, in non-human primates deep-layer SC neurons stimulated during a window of 120 ms initiate a saccade. For example, electrical stimulation of deep-layer SC neurons for 25 to 120 ms (e.g., 25, 50, 80, or 120 ms) results in a single saccade of the same size, as long as the stimulation duration remains ≤120 ms^23^. Doubling this temporal window (e.g., 240 ms) elicits two distinct saccades. This coding operation of SC neurons is known as a vector code, which distinguishes it from the firing-rate code of the BS nuclei neurons. The coding operation is important because consecutive saccades, independent of visual input but initiated by stimulation of deep SC neurons, appear to be dependent on this temporal coding scheme (120 ms). Crucially, this temporal window coincides with the length of a typical alpha cycle. The vector coding operation is not exclusive to deep-layer SC neurons and is found in other cortical areas such as in V1, parietal cortex, and medial frontal-eye fields ^19, 24^.

The existence of this coding operation raises the question how the cortex (e.g., V1) relates to the subcortical pathways involved in eye movement generation? SC neurons in deeper layers are controlled by cortical downflow independent of retinal input^17, 22^ and are predominantly innervated by complex cells in the deeper layers of V1^19^. Interestingly, alpha oscillations are stronger in these deeper V1 layers than in superficial layers^25, 26^. Electrical stimulation of deep-layer V1 neurons elicits saccades of constant size and direction, independent of the starting eye position, similar to the SC deep-layer neurons described above^19^. Moreover, cooling of V1 silences neuronal firing in deep SC layers but not in superficial SC layers^22^, and as a consequence suppresses saccade initiation. This cortical downflow regulation model of eye movements is further supported by the observation that right-hemisphere stroke, potentially interfering with this cortical downflow as in visual hemineglect patients, is associated with compromised eye movements of the contralateral left eye^27, 28^. In parallel to eye movement deficits, visual areas affected by brain lesions are associated with reduced spontaneous alpha activity relative to the non-affected hemisphere^29^. Thus, neurons in deeper layers of SC are innervated exclusively by the cortex, exhibit receptive fields similar to those elsewhere in the cortex, and follow a temporal coding scheme approximating the duration of an alpha cycle.

The present report provides empirical support for the hypothesis that alpha oscillatory activity is linked to oculomotor action. Based on the aforementioned putatively temporal relationship between cortical and subcortical circuits involved in saccade generation, the hypothesis prompts the following predictions. First, if power modulations of alpha activity are triggered by eye movements (e.g., saccades), alpha power should decrease following saccades. Second, this power modulation does not depend on visual input per se. Instead, it is independent of external input, as it reflects an internal mechanism. Thus, third, scalp topography of saccade-related alpha power modulations should be consistent with saccade direction. Fourth, the temporal window established by the alpha phase should manifest in an increased trial-to-trial phase locking prior to saccade onset. Finally, for reasons presented in the Discussion, the ecological validity of the theory should generalize across species.

Below we provide support for each of these predictions in a series of observations across species (human, non-human primate, insect), imaging modalities (EEG, ECOG, MEG), and cognitive states (rest, attention, and reliance on working memory). In summary, independent of visual input, alpha power is decreased following saccade onset and predicts saccade direction. The phase of alpha activity biases saccade onset and fixational tremor. Alpha power increase is related to a reduction of oculomotor action - a relationship evident during awake rest and WM tasks that generalizes to across species.

## Results

### Alpha power modulation follows saccade onset in humans and non-human primates

The first prediction about the relationship between saccades and alpha power is that alpha power should decrease (see^30^) after eye movement. In Experiment 1, 24 human participants performed a saccade calibration task. Participants were asked to follow a fixation cross as it was repositioned to the left, right, top, and bottom sections of the visual field. Saccade onsets were identified from the horizontal electrooculogram (EOG) and used to segment continuous EEG around saccade onset. Figure 1a illustrates the grand-average time-frequency representation of power (TFR) over occipital electrodes after the saccade relative to pre-saccade baseline. Cluster-permutation tests correcting for multiple comparisons (see Methods) revealed a reliable decrease in alpha (8-12 Hz) and beta (15-30 Hz) power after saccade onset, characterized by large effect sizes (Cohen’s d > 1, Figure 1a). Source reconstruction confirmed a distributed origin within parietal, motor and frontal brain regions (Figure 1b). Both the saccade-onset modulation of alpha-beta power (Figure 1c, cluster-permutation test, Cohen’s d > 1) and its cortical distribution (Figure 1d) were confirmed in non-human primate ECoG analyzed from a publicly available database (http://www.neurotycho.org/fixation-task). The strongest modulation of alpha power was evident in the vicinity of regions that have been reliably linked to saccadic control - frontal eye fields (FEF) and lateral-intra-parietal (LIP) area. However, it can be argued that a saccade inevitably leads to new input on the retina. In turn, the new retinal input could be the primary generator of the observed cortical power decrease rather than the saccade alone. This alternative explanation was examined in Experiment 2.

**Figure 1:**
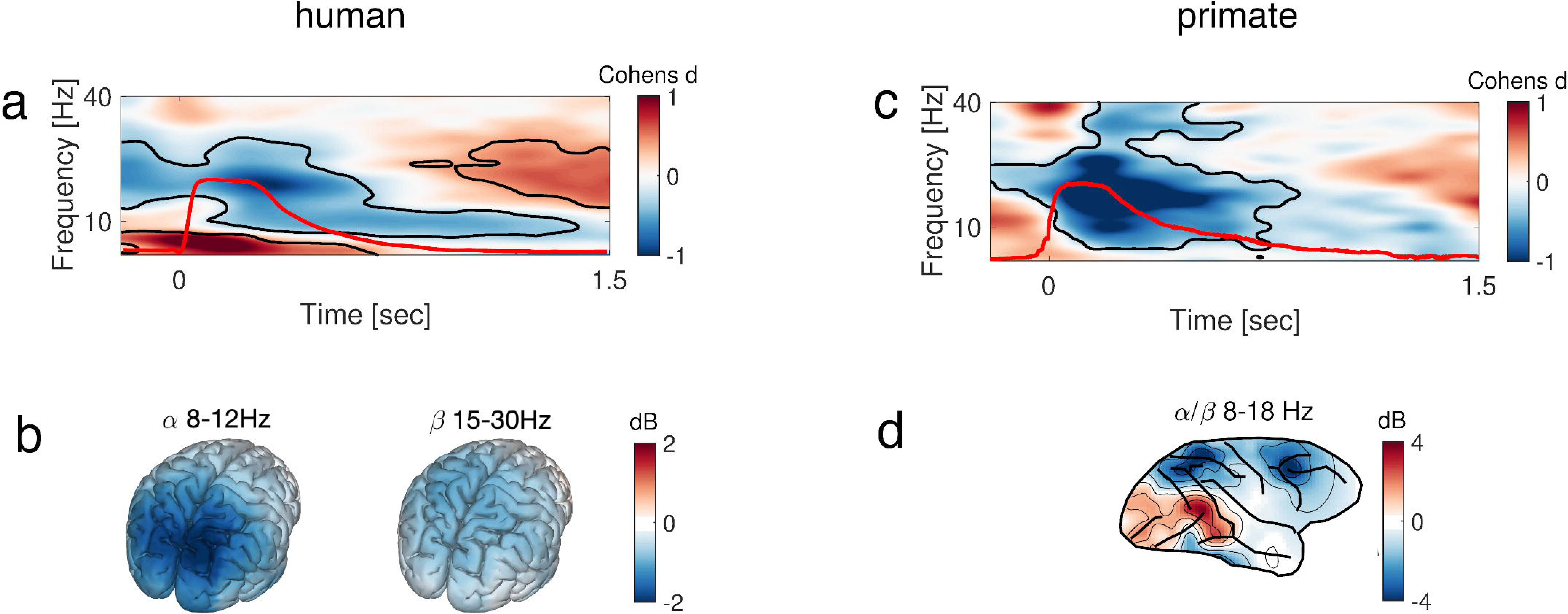
Alpha power modulation is induced by saccade onset. **a**- time-frequency representation of power (TFR) relative to pre-saccade baseline. Saccade onset is at 0 s, and the red line illustrates the grand-average time course of saccades derived from horizontal EOG. Color code denotes the difference from pre-saccade baseline (Cohen’s d). The black outline denotes clusters of significant condition differences (baseline vs. post-saccade, cluster permutation test corrected for multiple comparisons, p < 0.001). **b**- Source reconstruction of alpha/beta power modulation. **c**- Same as (a) but for non-human primate. The red line illustrates the grand-average time course of saccades derived from an eye-tracking device. Color code as in (a). **d**- Topographical distribution of alpha/beta power modulation across all ECoG electrodes in non-human primate. Key structures of the fronto-parietal attention network such as FEF and parietal cortex are consistent across both species.

### Alpha power modulation is independent of retinal input

In Experiment 2, the dependence of alpha modulation on retinal input was tested by initiating saccades performed with closed eyelids, thus excluding retinal input. Three volunteers were asked to perform spontaneous saccades with eyes closed while saccade time course was monitored by EOG. Alpha power was reliably modulated following saccades (Figure 2a, cluster-permutation test, Cohen’s d > 1), essentially replicating the results in Figure 1a. However, this finding does not necessarily translate to a covert attention scenario, where an external cue typically prompts the power modulation. An example is the delayed-saccade-to-target (Posner) task (Figure 2b), which is known to elicit reliable modulation of alpha activity with eyes open yet in the absence of visual stimulation^31, 32^ - covert attention. The literature on covert attention has established that alpha power modulation is lateralized according to the maintained position^11, 33^ (confirmed in Figure 2c). That is, maintenance of previously cued position in the left visual field is associated with contralateral right-hemisphere suppression of alpha power and vice versa (thus, place topography). It has been suggested that position maintenance during the delay interval, despite any visual input from the computer screen, is nonetheless associated with saccades toward the previously cued location ^12, 13^. As these experiments are typically performed under normal ambient light conditions, new retinal input ultimately follows the saccade. Consequently, this retinal input might elicit an evoked response followed by modulation of alpha power. Eliminating the effects of light also eliminates retinal responses prior and post saccades. This was pursued in Experiment 3.

**Figure 2:**
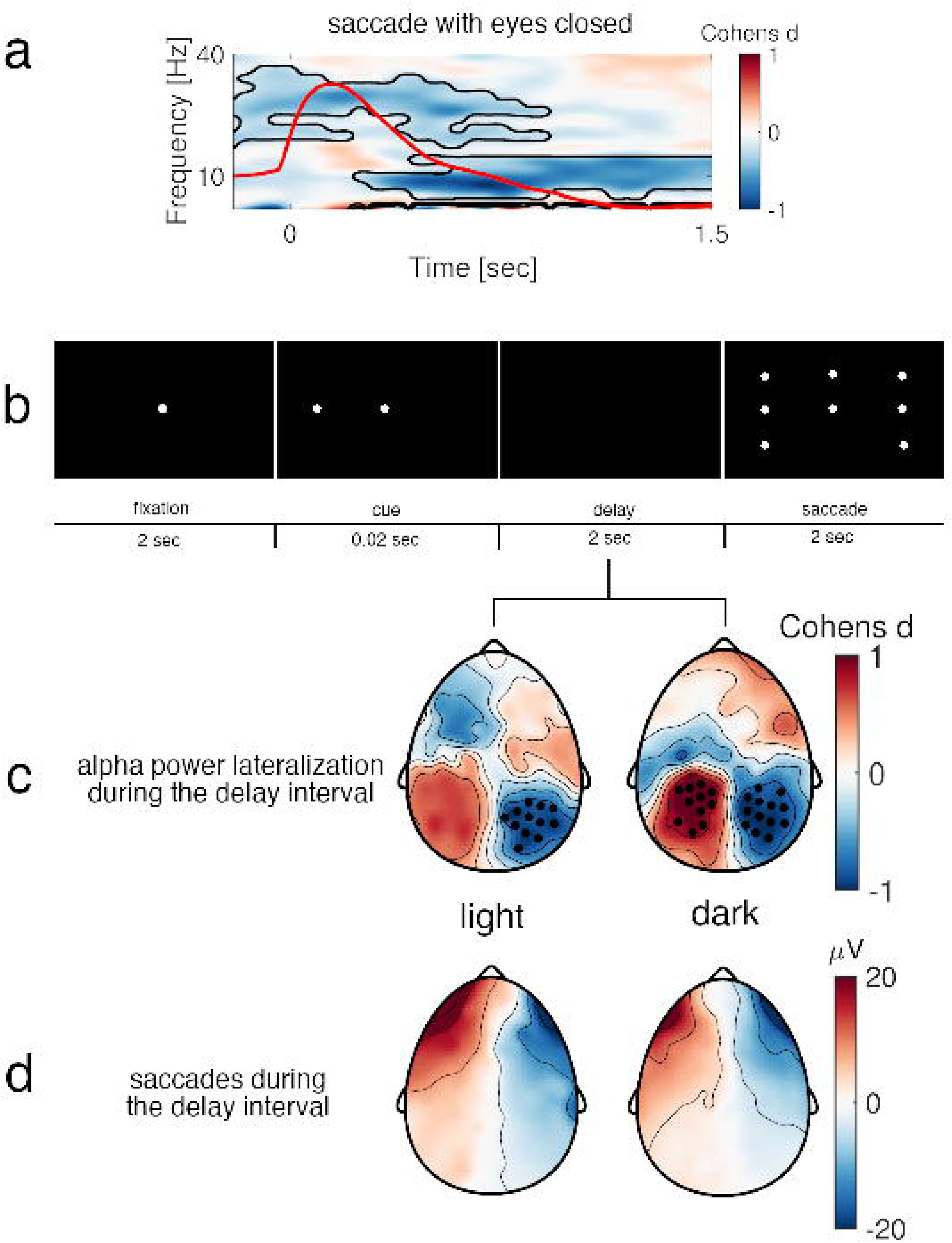
Saccade-induced alpha power modulation is independent of retinal input. **a**- Alpha power modulation with respect to pre-saccade baseline. Saccades were performed with eyes closed. The red line illustrates the grand-average time course of saccades derived from horizontal EOG. The black outline denotes clusters of significant condition differences (baseline vs. post-saccade, cluster permutation test, p < 0.001). Color code denotes the size of the difference from pre-saccade baseline (Cohen’s d). **b**- Experimental design illustrating the Posner delayed-response saccade task for both light and dark conditions. In this example the white dot in the left visual field indicates that the participant should saccade to that location after the delay interval. **c**- Maintenance of cued position prompts alpha power lateralization (right-hemisphere = contralateral alpha suppression: place topography) during the delay interval. Place topographies reflecting alpha for left minus right cued position are consistent across regular ambient light conditions (left) and full darkness (right). Black dots denote electrodes in clusters showing significant condition differences (cluster-permutation test, p < 0.05). Color code denotes the size of the left minus right cued location difference (Cohen’s d). **d**- Scalp topography of the left minus right cued position difference during the delay interval computed from the raw EEG data without ICA artifact control. A prototypical saccade topography is evident in both conditions: during normal ambient light (left) and full darkness (right).

In the present Posner task (Experiment 3), a cue is presented for 20 ms at one of seven possible locations on a visual display. Participants were instructed to maintain (covertly attend to) the position of the cue during a 2 s delay interval (while asked to maintain their gaze on a central fixation position), after which they executed a saccade in the direction of the maintained position. Analyses focused on the delay interval. Change in gaze position (e.g., following saccade) during full darkness is possible and likely. Crucially, however, in the absence of light there is no input-related activity either on the retinal level or the cortical level. Thus, if indeed saccades are present during the delay interval and are associated with cortical alpha power modulation independent of retinal input, this should be the case during both ambient light and full-darkness conditions. Hence, the experiment was performed twice: once under normal ambient light conditions and once in darkness (implemented by designing an aluminum grid with 8 LEDs, each of which could be triggered individually, ensuring a state of full darkness in the recording room during the delay interval).

Alpha power suppression contralateral to attended visual field position (thus, place topography) was present during both experimental conditions (Figure 2c, cluster-permutation test, effect size Cohen’s d > 1). This consistency across conditions indicates that modulation of alpha power cannot be explained by retinal input, because, in the absence of light, changes in gaze position following saccades do not elicit any external input-related retinal and subsequent brain activity. A common feature of studies utilizing posterior alpha power lateralization during covert attention, including the present Posner task, is the correction of saccades and eye blinks by means of ICA. Because the rotation of the eyes in orbit as well as the eye muscles supporting the rotation produce characteristic EEG scalp topographies, independent component analysis (ICA) is an efficient approach for removing these topographies prior to further analyses. However, upon every movement in the human body a copy of the signal sent to the muscles is also sent to the brain, commonly termed a corollary discharge. It is noteworthy that ICA correction eliminates the scalp topography and time course of the eye movements as a result of eye muscle contraction and the rotation on the eyeball. Yet the potential consequence of eye movements for visual cortical processing including corollary discharge signals could persist. In light of previous evidence on saccadic activity during the delay interval^13^, we re-evaluated the raw, non-ICA-corrected data during both conditions to examine the presence of saccadic eye movement topographies during the delay interval. If saccades are present and consistent with the cued location, saccade-typical scalp topography toward left cues will be opposite to saccade-typical scalp topographies toward right cues. Consequently, the difference in saccade topography (left minus right cue) will also result in saccade-typical scalp topography. Indeed, that is what we found. Figure 2d illustrates the difference in non-ICA-corrected EEG scalp topography of left vs. right cued location during the delay interval. This result confirms that 1) saccades are performed even during full darkness and 2) their direction is consistent with the maintained cued location.

In sum, given (1) reduced alpha power following saccade onset, (2) even during saccades with eyes closed, and (3) independent of retinal input, yet consistent with saccade direction, we conclude that place topographies emerge in line with the direction of a previously performed saccade. Thus, if saccades are associated with the instantiation of the place topography during the delay interval in the Posner task (Figure 2c), lateralized saccade direction should be associated with lateralized modulation of alpha power. Using a state-of-the-art eye-tracking device and simultaneous EEG acquisition, a further validation of this hypothesis was pursued next.

### Lateralized saccade direction elicits lateralization of posterior alpha power

In Experiment 4, human volunteers (N = 143) performed a prosaccade task in two sessions, separated by a week^34^. Eye-tracking was used to identify and segment the data according to saccade onsets. Figure 3a illustrates that lateralized saccade direction (left minus right) is associated with lateralized modulation of alpha power (cluster-permutation test, effect size Cohen’s d >1). This modulation inverts following the opposite saccade toward the fixation point (i.e., alpha lateralization is in the opposite direction when participant makes a return saccade) and provides further evidence that place topographies, i.e. the topography that allows inference about saccade direction based on alpha power scalp distribution, can be elicited as a consequence of oculomotor action. The topography is reproducible at retest assessment (Figure 3b), with fair test-retest reliability as assessed by the interclass correlation (ICC = 0.6, Figure 3c). Of note, these results remain essentially the same (see supplemental Figure S1) after controlling for the potential confounding influence of the ocular activity using recent advances in modeling of combined eye-tracking and EEG data^35, 36^. Hence, place topographies emerge following saccade initiation and are consistent with saccade direction. Furthermore, as suggested by the results of Experiments 2 and 3, place topographies can be generated endogenously, that is, independent of retinal input during full darkness. The overlap between the temporal window of the SC in saccade generation and the length of an alpha oscillatory cycle (see Introduction) introduces the fourth prediction, that the temporal window established by the alpha phase should manifest in increased trial-to-trial phase coherence prior to saccade onset. This hypothesis was evaluated next.

**Figure 3:**
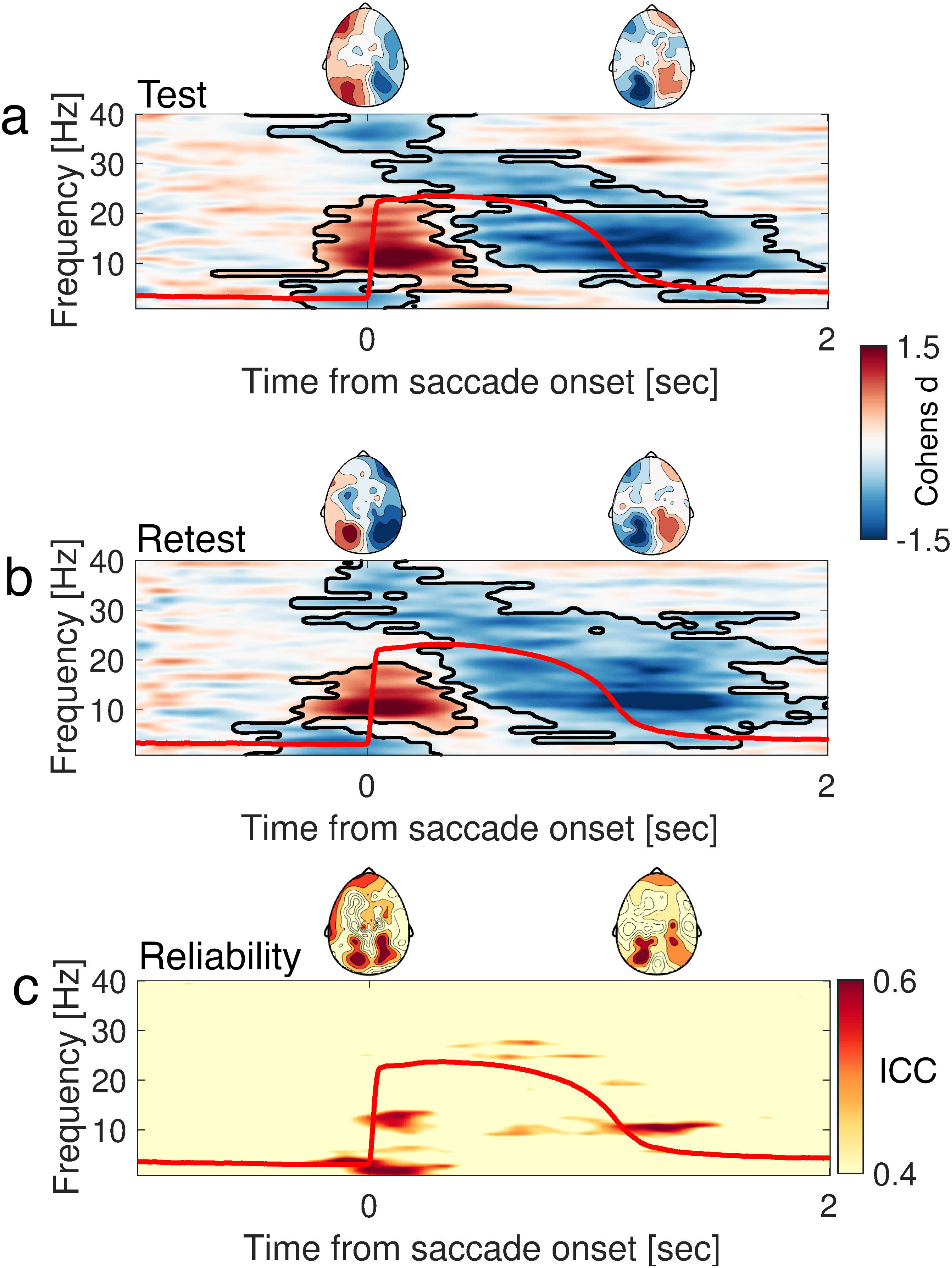
Place topographies are consistent with saccade direction. **a**- Time-frequency representation of power (TFR) derived from the difference (LEFT HEMISPHERE_(left-right saccade)_- RIGHT HEMISPHERE_(left-right saccade)_) averaged over occipital electrodes. Warm colors represent an increase and cool colors a decrease of oscillatory power. Black outlines illustrate clusters of activity confirming significant condition differences (cluster-premutation test, p< 0.001). Place topographies illustrate the spatial distribution of alpha power around prosaccade onset −100 to 100 ms and 700-900 ms reverse saccade onset toward fixation respectively. The red line illustrates the grand-average time course of saccades derived from the eye-tracking device. **b**- Same as in (a) but for the retest examination of the same participants a week later. **c**- Time-frequency representation of reliability. Warm colors indicate an increase in interclass correlation (ICC).

### Saccade onset is phase-locked to the phase of alpha activity

Previous research^37–40^ confirmed an increase in inter-trial phase coherence (ITC) of alpha activity prior to saccade onset. Hence, we analyzed ITC using the data from Experiment 4 above to examine the relationship between the alpha phase and saccade generation. ITC of each of participant was computed on the time series locked to all saccade onsets identified by the eye-tracking device (number of trials in test: M/STD = 1080/310, range = 589-893, retest: M/STD = 1052/301, range = 60-1852). The analysis used a sliding window approach. Therefore, pre-saccade effects were evaluated up to −250 ms prior to saccade onset to prevent contamination of the analysis window (500 ms length) by post-saccade activity. Analyses were carried out in source space (see Methods). The presence of ITC is typically evaluated as contrasted against surrogate data. This contrast is needed, as ITC can be observed by chance. Circularly shifting the time series several hundred times, thus destroying the inter-trial relationship, and re-computing the ITC, generates surrogate data. This contrast addresses the presence or absence of ITC in the data, which as mentioned above has been reported in numerous studies. To asses, not only whether ITC emerged simply by chance but also how reliable it is, we assessed the consistency across acquisition time points rather than the mere presence tested against surrogate data. Figure 4a illustrates the test-retest reliability of alpha phase coherence prior to saccade onset. Consistent, albeit with somewhat modest reliability (ICC =.4), alpha phase coherence prior to saccade onset was confirmed in bilateral FEF and orbitofrontal cortex (OFC). Following saccade onset, alpha phase coherence was sustained in primary visual cortex over ∼1 s, with fair reliability ICC = .6 (Figure 4c). Note that during this period after saccade onset the power of alpha activity is decreased. Contrary to the general assumption that alpha power decreases signal desynchronized brain activity, the phase of alpha activity is still functional in the sense that it enables a frequency specific trial-to-trial coherence in primary visual cortex. The coincidence between the length of a typical alpha cycle and the temporal coding scheme for saccade generation in the SC indicates a relationship between the ongoing alpha phase and saccade occurrence.

**Figure 4:**
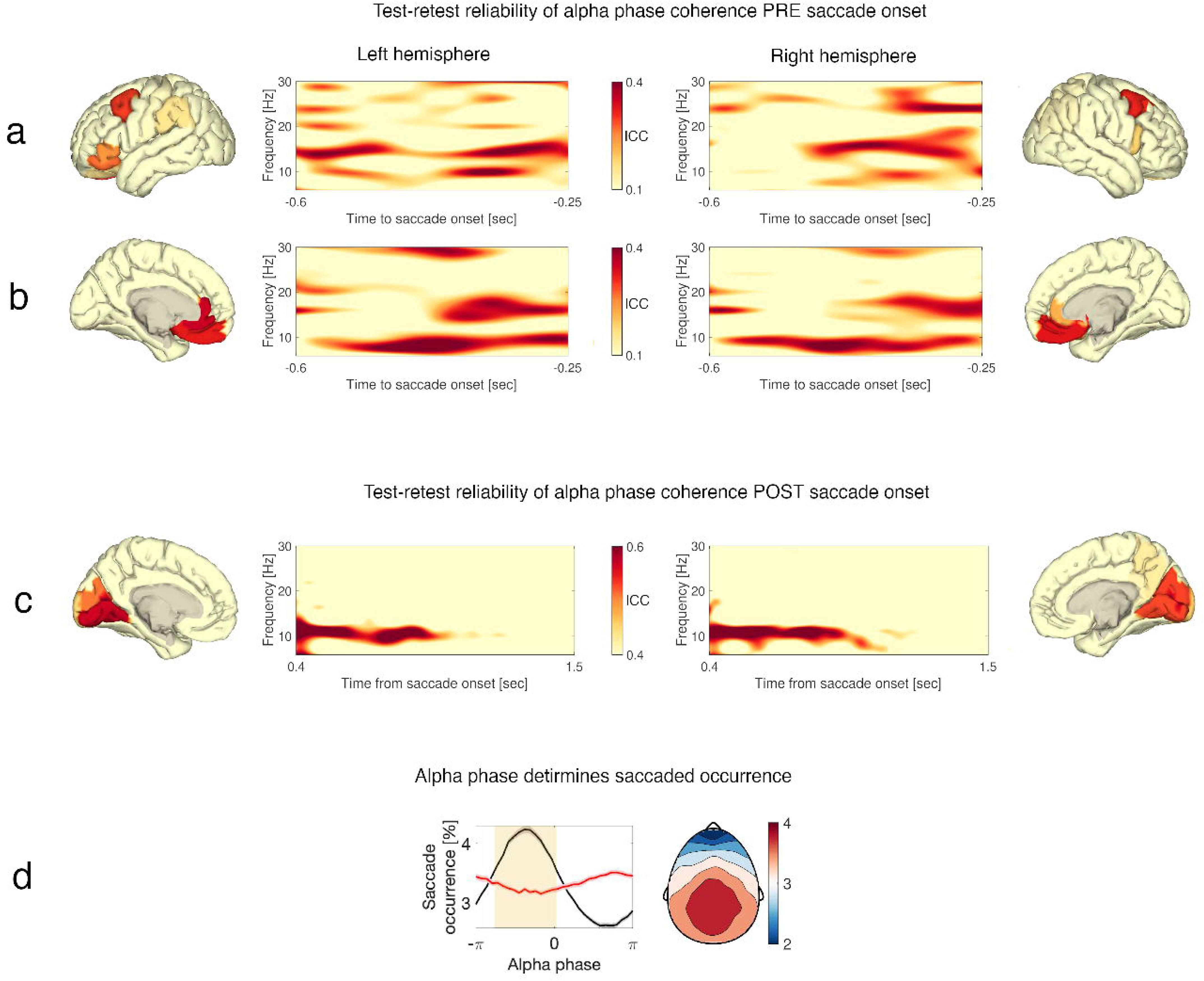
Saccade occurrence is locked to alpha phase. **a-b** Time-frequency representation of test-retest reliability in inter-trial phase coherence (ITC) prior to saccade onset. Warm colors indicate higher ICC values. Source reconstruction revealed that the highest reliability in ITC values were found within the FEF and OFC. Time-frequency spectrograms correspond to these highlighted parcels. **c**- Same as (a) and (b) but showing test-retest reliability in alpha phase ITC after saccade onset. After initial broadband increase in ITC due to evoked activity, a sustained effect in alpha phase coherence is observed. **d**- Saccade occurrence as a function of alpha phase averaged across participants for the original data (black line with SEM) and surrogate data (red line with SEM). Cluster-based permutation testing revealed significant group differences between original and surrogate data with clear posterior topography illustrated for the cluster denoted by the yellow shading.

Next we evaluated the extent to which saccades occur at a particular alpha phase. For each participant, a Hilbert transformation using an 8-12 Hz band-pass filter was applied to the continuous EEG. Thus, each sample points during saccade occurrence was also assigned an alpha phase value in radians. For each electrode and participant, saccade occurrence (number of saccades in % per phase bin from total number saccades) was determined for 30 phase bins distributed from -π to π. Figure 4d illustrates a clear phase preference for saccade occurrence as a function of alpha phase. This effect was characterized by a posterior topography and differed significantly from surrogate data (cluster-based permutation test, effect size Cohen’s d > 1). The latter was obtained after circularly shifting the continuous time series (± 100 ms) in each one of 100 permutations, and a subsequent circular average across permutations (e.g., red line in Figure 4d).

Taken together, results confirm that saccade onset is influenced by the cortical alpha phase and that post-saccade alpha power modulation reflects place topography consistent with saccade direction. The latter conclusion is based so far on experiments 2 and 3, where saccades were directed toward left or right hemifield only. As the literature has established a link between alpha power modulation and attention direction^8–10, 15^ and similarly between attention direction and saccades^14, 41, 42^, next we addressed to what extent this same representation occurs for saccades generated omnidirectionally during awake resting conditions in the absence of any task context.

### Visual space is represented by spontaneously generated and spatially specific place topographies

During awake resting states, saccades are produced endogenously and omnidirectionally, covering the entire visual field. If alpha power is related to oculomotor action, it is conceivable that place topographies exhibit spatial distributions beyond just left vs. right hemispace. The discovery of place cells in the rodent hippocampus constitutes one of the more robust and well established findings in neuroscience^43^. Briefly, as the animal explores the environment, sets of neuronal cells fire sequentially in accordance with its current spatial position. This phenomenon has been confirmed not only during active exploration but as a self-organized pattern, independent of environmental cues^44–46^. The path navigated by the animal can be represented by a set of tuning functions describing the firing (or tuning) of an assembly of neurons as a function of position within the path (e.g., Figure 6F in ^44^). Thus, the sum of the tuning functions is a model approximation of the physical path length navigated by the animal.

Similar observations have been made in human EEG investigating visual^8–10^ and auditory^15^ spatial attention. Specifically, it has been shown that the entire visual field or auditory space around the listening individual (e.g., circular path length of 0-360°) can be modeled by a set of tuning functions per spatial location. A common finding in these studies is that, independent of sensory modality (visual or auditory cue), the modulation of alpha power in visual brain areas reveals spatial tuning to a target location. This is utilized, at least in part, by oculomotor (e.g., saccade) activity^12–15^. Crucially, the saccadic spike potentials evoked by spontaneous saccades during free viewing 1) are constrained by the field of action of the ocular muscles^47^ (see below, Figure 5a) and 2) exhibit “retinotopically” organized scalp topographies^41^. Bridging rodent and human electrophysiology, we reasoned that, conceptually, the endogenous (i.e., independent of external cues) nature of the tuning patterns in the rodent could also apply to place topographies in humans. That is, place topographies are not only elicited via visual cues but generated spontaneously as a consequence of the omnidirectional distribution of saccades.

**Figure 5:**
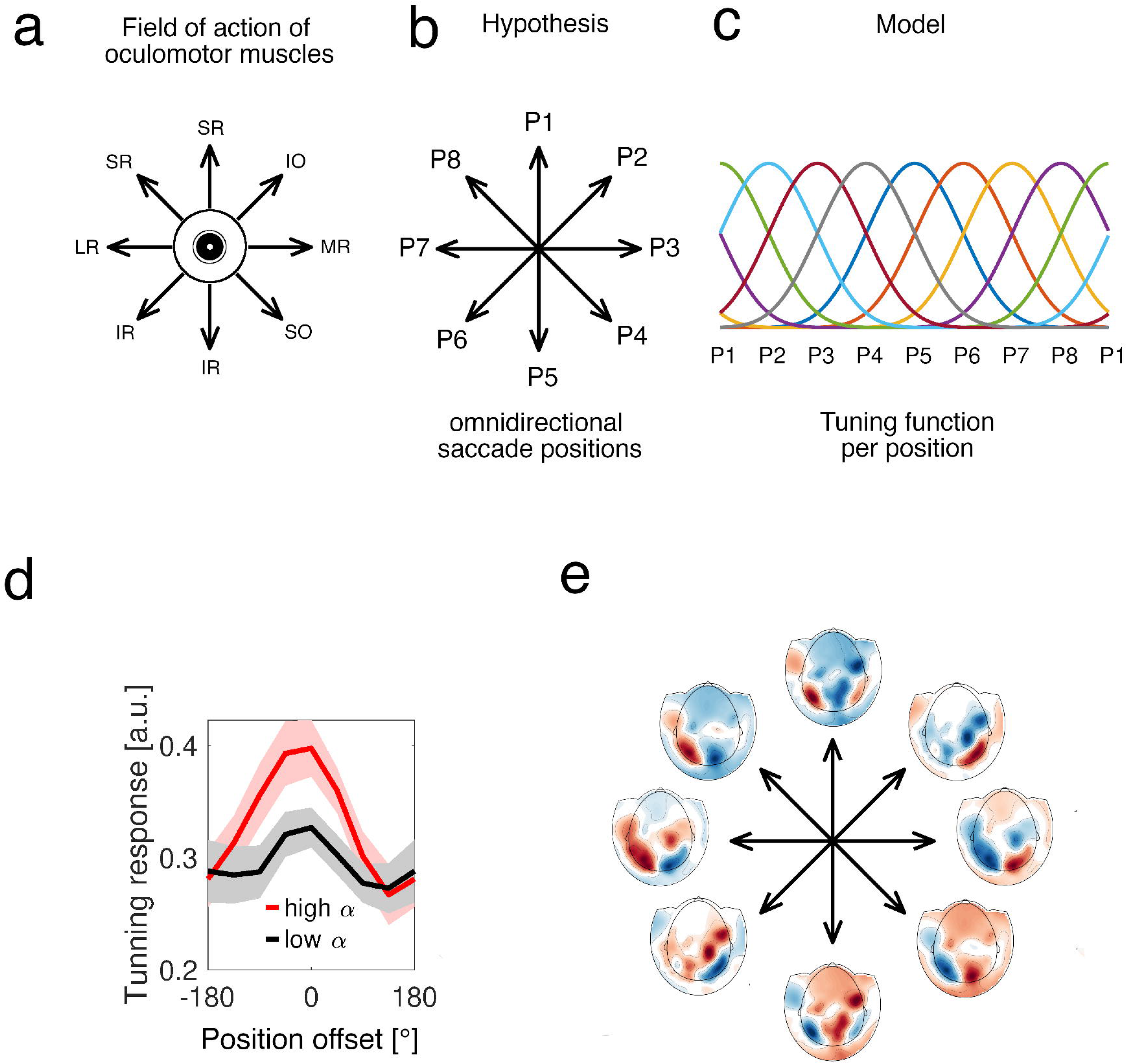
Spontaneous alpha power fluctuations reveal omnidirectional saccade-based exploration. **a**- During awake resting conditions, saccades are generated spontaneously within the boundaries of the field of action of the ocular muscles covering the entire circular visual space: superior rectus (SR), inferior oblique (IO), medial rectus (MR), superior oblique (SO), inferior rectus (IR), lateral rectus (LR). **b**- Hypothetically, each saccade direction can be assigned a position (P1 to P8). **c**- A model of hypothetical tuning functions that spans the entire position space. Each tuning function has high sensitivity for a particular position. **d**- The tuning response derived from the spatial weights W and the test data approximates a bell-shape function. The abscissa denotes the position offset in degrees (°) from a common center. The tuning response (ordinate) is maximal at 0°, confirming the alignment of the hypothetical tuning model with the actual data. The tuning response is stronger for data dominated by high (red) than by low (black) alpha levels of power. Error shading denotes 1 SEM. **e**- The spatial distribution of the tuning weights (W) approximates ‘retinotopic’ organization. Warm and cold colors denote the variation in weight magnitude over posterior sensor sites as the most informative regions.

To test this hypothesis, in Experiment 5, a publicly available MEG dataset was used (https://www.humanconnectome.org/study/hcp-young-adult/project-protocol/meg-eeg). A forward-encoding model^48, 49^ was applied to resting-state eyes-open recordings. During data acquisition, participants are asked to maintain central fixation and avoid eye movements. The presence of saccades, at minimum along the horizontal (left/right) axis, is a prerequisite for the postulated hypothesis. Thus, using horizontal EOG, the indices of left and right saccades were identified first. Supplemental Figure S2 confirms that saccades along the principal left-right direction were indeed present in the data, despite the instruction to maintain central fixation. Furthermore, alpha power was modulated after saccade onset consistent with the observations in Experiments 1-4. Next, the ongoing data were randomly segmented into epochs of 1 s. Each epoch was assigned a random position (P1-P8, Figure 5b), and the forward-encoding model was applied. This procedure was repeated 1000 times, and the tuning response was averaged over these repetitions. The position assignment was chosen randomly, but the positions themselves were not. Rotation of the eye is achieved by the primary field of action of the eye muscles (Figure 5a). Research has identified eight directions of gaze for which an individual eye muscle is the primary actuator of the eye in orbit^47^. In a clock analogy and considering the right eye, the lateral and middle recti correspond to rotation toward 9 (lateral) and 3 o’clock (nasal), the inferior and superior obliques to 1:30 and 4:30 o’clock (nasal), the superior and inferior recti to 12 and 6 o’clock (up/down), and the superior and inferior recti toward 7:30 and 10:30 o’clock (up and lateral vs. down and lateral). Accordingly, the hypothesized positions were chosen in register with the field of action of the extraocular muscles (Figure 5a): P1 and P8- superior rectus (SR), P2- inferior oblique (IO), P3- medial rectus (MR), P4- superior oblique (SO), P5 and P6- inferior rectus (IR), P7- lateral rectus (LR). Amongst the realm of possibilities of combined contraction and relaxation of multiple muscles enabling the entire faculty of eye movements, these positions have been identified as the positions where one muscle provides the dominant force^47^. Similar to rodent model, the sum of the tuning functions per position is a model approximation of the visual field covered by eye movements (Figure 5c).

Under the null hypothesis of no associations between assigned position and MEG data, one would expect a tuning response characterized by a flat line instead of gaussian shaped response (Figure 5d). An affirmative test for this hypothesis would have involved replacing each epoch’s data with random noise. However, this procedure would also destroy the cyclostationarity in the data, yielding a somewhat biased comparison. Instead, we divided the 1 s epochs into low and high alpha power based on a median split and performed the forward-encoding modeling as described above. If there were no influence of alpha power on spatial decoding performance, one would expect no differences in the tuning response between high and low alpha power data segments.

The present analysis rejected this hypothesis (Figure 5d). The tuning response was higher in trials dominated by high alpha power (p < 0.001, cluster-based permutation test, Cohen’s d > 1), and its bell shape confirmed the hypothesized specificity. The topographic charting of the weights used to fit the model to the data displayed a characteristic pattern resembling a “retinotopic” map (Figure 5e). Finally, these place topographies were circularly shifted in accordance with the characteristic lateralized pattern specific to the azimuth (left-right) direction (e.g., Figures 2c and 3a,b). This pattern confirms the presence of endogenous place topographies informing the brain about its self-initiated direction of oculomotor action.

To summarize, the results from Experiments 1 to 5 provide empirical support for the four predictions derived from the oculomotor hypothesis outlined in the Introduction:

1. power modulation of alpha activity relates to eye movements (Experiments 1-4)
2. this power modulation does not depend on the visual input (Experiment 2-3)
3. scalp topography of saccade-related alpha power modulations is consistent with saccade direction (Experiment 4-5)
4. the temporal window established by alpha phase manifests in an increased trial-to-trial phase locking around saccade onset (Experiment 4).

Alpha power is utilized as a dependent variable across a variety of cognitive tasks, most frequently in the study of working memory (WM). What is the relevance of the proposed relationship between oculomotor action and alpha power modulations during actual task conditions? This question is pursued in the next section.

### Linking alpha oscillations to working memory through oculomotor action

From early on, alpha power modulation has been associated with cognitive functions such as attention and working memory (WM). A well-established observation is that the maintenance of several items in traditional WM tasks (Sternberg or N-back) is associated with a power modulation in posterior alpha activity^50–52^. For example, alpha power is found strongest for e.g. 7 items, lower for 4 items and lowest for 1 item. In contrast, the same WM construct studied in the N-back task (e.g., matching a current item to 2-items back) is associated with the opposite pattern: lower alpha power in the higher WM load conditions, such as 2-back vs. 1-back or 0-back conditions^53–55^. That is, depending on the specific task, high WM load is associated either with high (Sternberg) or low (N-back) levels of alpha power. It has been suggested that this contradiction can be solved by assuming that it is the sensory load that drives the modulation of posterior alpha power^56^ (i.e., simultaneous encoding and maintenance in N-back tasks, but sequential intervals of encoding and maintenance in Sternberg tasks). In this section we demonstrate that this contradiction can be resolved by the oculomotor hypothesis proposed above. In particular, the oculomotor action is inversely related to power modulations in the context of WM tasks.

In Experiment 6, we analyzed EEG from an individual performing a Sternberg WM task from a publicly available dataset^57^ as well as the N-back dataset provided by the human connectome consortium (https://www.humanconnectome.org). In the Sternberg task, EEG was recorded during the maintenance of 1, 4, or 7 items in working memory. Neither dataset provides eye tracking data. As an alternative, time courses of oculomotor activity were extracted from the continuous data after performing an ICA and selecting only those components judged as reflecting eye movement activity. Subsequently, two electrodes near the left and right outer canthi were selected and converted to a bipolar derivation, resulting in a horizontal EOG time series. Oculomotor activity (e.g. muscle contraction) typically manifests as a broadband increase in the power spectrum^58^. If oculomotor activity is captured by the hEOG one could compare its variation with WM load together with the occipital modulation of alpha power. Thus, we compared power spectra over occipital electrodes with the hEOG spectra. As illustrated in Figure 6a, higher WM load was associated with longer reaction time and a concomitant higher occipital alpha power. Crucially, the broadband power (10-200Hz) reflective of oculomotor activity followed the opposite pattern - the higher the WM load, the lower the oculomotor activity. The contradictory modulation of alpha power with WM load between Sternberg and N-back task was confirmed (Figure 6a vs. 6b), whereas the oculomotor-alpha power relationship was preserved in the same direction (Figure 6b). This confirms the above proposition that the oculomotor action modulated alpha power rather than WM load per se.

**Figure 6.**
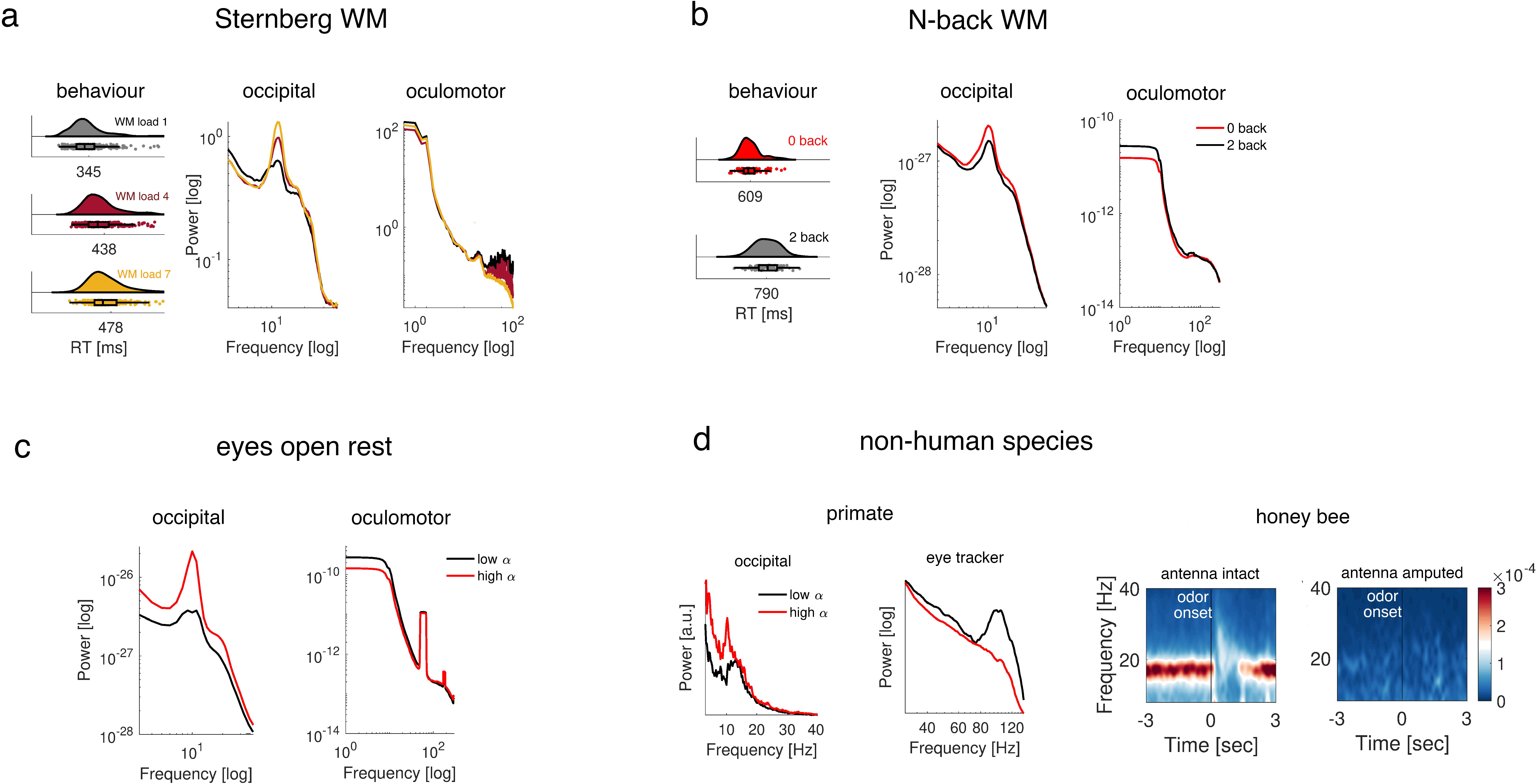
High alpha power is associated with reduction in oculomotor activity across tasks and species. **a**- Representative participant performing a Sternberg working memory task. The data are analyzed during a 3 s maintenance interval without visual input. Higher WM load is associated with longer reaction time (histogram of RT across trials, left) and higher occipital alpha power (middle). Oculomotor activity is inversely related to alpha power (right): higher oculomotor activity is associated with lower alpha power. Oculomotor activity is derived from the hEOG channel, and the typical increase in broadband activity resulting from muscle contraction is illustrated on a log-log scale (right). **b**- Group analysis across 83 participants from the human connectome project performing an N-back WM task (histogram of RT across participants, left) confirms the association described in (a): lower alpha power during the higher WM load condition (2-back) is associated with higher oculomotor activity. **c**- The same association is present within participants during the awake resting state condition. On an intraindividual level, high alpha power (red) is associated with lower oculomotor activity and vice versa. **d**- This association is also evident in the non-human primate. The two left panels in (d) show a clear association between a high level of high-frequency saccadic eye movements and low occipital alpha power. In the honey bee (d), the jerky movements of the sensory antennae serve a function similar to that of saccadic behavior in humans and non-human primates. Honey bees have a dominant rhythm that is spontaneously generated and modulated upon odor presentation (0 ms). Electrophysiological recordings in the same animals (N=5) before (left) and after (right) amputation of the sensory antenna confirms the link between sensory-motor action and the dominant rhythm (reprinted with permission from^59^).

### The association of oculomotor action and alpha oscillations generalizes across states and species

This link between oculomotor activity and alpha power was not specific to memory load but generalized to awake resting state in humans (Figure 6c) as well as non-human species (Figure 6d). Awake resting-state recordings displayed higher oculomotor activity when alpha power was lower (Figure 6c). Similar analysis of the primate data revealed more high-frequency variability in the eye-tracking recording during low alpha power (Figure 6d). This finding generalizes to the invertebrate brain, as we have demonstrated previously^59^. The “jerky” movement of the sensory antenna in the honey bee has utility similar to that of eye movements in primates - active sampling of the environment. In addition, insect flight pattern, termed “body saccades”, has been described as a means by which gaze patterns are stabilized^60, 61^. Honey bees show a neural rhythm comparable to alpha oscillations in humans^59^, in that the rhythm is spontaneously generated and attenuated following sensory stimulation (Figure 6d). Eliminating the sensorimotor action of the sensory antennae (i.e. amputation of the sensory antenna) eliminates the rhythm.

Given that oculomotor action is not limited to saccades, a further question is whether the alpha/oculomotor relationship substantiated by the results presented above generalizes beyond saccades. As outlined in the Introduction, miniature eye movements conceived as a fixational tremor are produced continuously. Their manifestation is evident in the high-frequency activity of the simultaneous EOG or eye-tracking timeseries and is present during awake rest and cognitive tasks. Hence, is the alpha/oculomotor relationship continued also during such miniature eye movements? An exploratory analysis was carried out to address this question.

### Alpha phase modulates fixational tremor

An exploratory analysis of a single volunteer served to highlight the relevance of alpha power and alpha phase in oculomotor action. Figure 7a illustrates the ongoing EEG signal (low-pass filtered at 30 Hz) superimposed on the horizontal eye-tracker time series while the participant engaged in the prosaccade task. The spindles of alpha activity with high amplitude are present during fixation intervals, which per design correspond to the pre-saccade baseline (e.g., fixation) period utilized in the group analysis in Figure 3. A closer look at one of these high-amplitude spindles (Figure 7b) confirmed previous reports of continued eye movements in the form of fixational tremor. Even within a temporal frame of one alpha cycle, the degree of horizontal eccentricity of the fixational tremor is visibly modulated by the phase of the ongoing alpha activity (Figure 7c). The left-right deviation from fixation is largest at the trough and attenuated at the peak of the alpha cycle. Cross-frequency-coupling analysis performed on the entire 5 min recording confirmed that alpha phase modulates the high-frequency amplitude variation in eye movement (Figure 7d). This cross-frequency relationship between cortical oscillations and eye-tracking activity was consistent across acquisition time points (e.g., test and retest of the same participants, Figure 7d). That is, the group-level results demonstrating the modulation of alpha power around the saccade onset (Figure 1) and the relevance of alpha phase prior to saccades, i.e. during fixation, could be reproduced on the single individual as well as on the group level.

**Figure 7:**
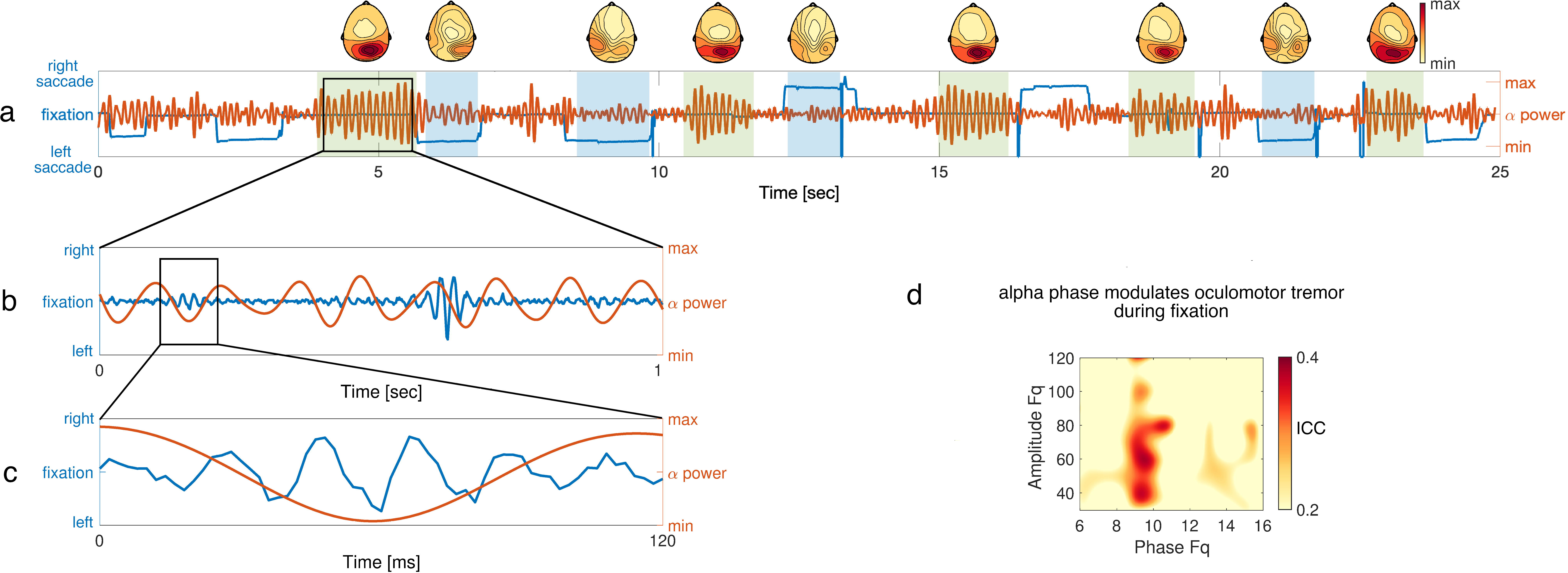
Alpha power and phase associated with oculomotor control. **a**- Raw EEG trace (red) and simultaneous eye-tracking horizontal time course (blue) during the prosaccade task. Scalp topographies of alpha power are averaged across the time intervals highlighted by the green (during fixation) and blue (nonfixation) shading. **b**- Same as (a) but for the time window highlighted by the black rectangle in (a) and the eye-tracking time course high-pass filtered at 30 Hz. **c**- Same as (b) but for the time window of a single alpha cycle highlighted by the black rectangle in (b). **d**- Cross-frequency coupling between the alpha phase (abscissa) and the amplitude of eye-movement variability in the eye-tracking data (ordinate). Warm colors indicate interclass correlation coefficient (ICC) quantifying test-retest reliability across all participants.

Present analyses support the overall conclusion that alpha oscillations are closely linked to the variability in oculomotor activity during tasks and rest. Alpha power is attenuated following saccades, independent of external input (Experiments 1-4). This power modulation is consistent with saccade direction manifesting in place topographies during task (Experiments 2-4) and rest (Experiment 5). Saccade occurrence co-varies with the ongoing alpha phase (Experiment 4). This link between alpha activity and oculomotor action provides an explanation for the previously reported, apparently contradictory association between alpha power and WM load in the context of Sternberg vs. N-back WM tasks (Experiment 6). Periods of fixation, characterized by reduced oculomotor action, were also associated with increased alpha power (Experiment 4 & 6). Finally, during fixation, eye movements do not simply stop but segue into fixational tremor. Similar to saccades, this tremor varies with the phase of the ongoing alpha activity (Experiment 4, Figure 7). The variety of findings presented here support the contention that visual alpha activity reflects the implementation of an oculomotor action command that is distributed across visual cortical areas in order to inform the brain about the likelihood of impending input - a corollary discharge.

## Discussion

Alpha oscillations are commonly associated with human cognitive processes such as attention^62^, memory^63, 64^, and language^65, 66^, yet the functional specificity of the rhythm and the mechanism by which it relates to cognitive function is to date unclear. The present summary of studies spanning species, cognitive operations, states, and methods advocates for a new oculomotor theory linking alpha oscillations to oculomotor control. Based on general principles that vision is defined thru eye movements and bodily movement is controlled by the central nervous system, the theory explains why power modulations of alpha oscillations are observed in nearly every cognitive construct studied to date. This is because cognitive operations entail coordinated eye movements and eye movement control depends on cortical alpha oscillations. Absence of coordinated eye movements is also characterized with absence of alpha activity (e.g. as in neonates^4, 5^, congenital blindness^67, 68^ or locked-in state^6, 7^). Saccade occurrence and fixational tremor are influenced by the phase of alpha activity, whereas alpha power, high during intervals of fixation, is reduced following a saccade execution. Across states (e.g., task and awake rest), cognitive operations (overt and covert attention, WM), and species (human, nonhuman primate, and insect), high oculomotor activity is associated with low alpha power, and high alpha power is a signature of reduction in oculomotor action. Can this redefinition of the functional significance of alpha oscillations be assimilated with common concepts on their role in cognition?

### Alpha oscillations link to cognition thru their functional role in eye movement control

Evidence that alpha power modulation signals directed attention^69^, increases with working memory load (e.g. ^50^), and facilitates successful memory retrieval (e.g. ^70, 71^) proved Berger’s original association (and functional definition) of alpha as representing an idling brain state wrong. Alpha power increase with executive demand encouraged to define the role of alpha as top-down or inhibitory control ^71^. Different levels of alpha power reveal the active routing of information throughout the brain by inhibiting task-irrelevant brain regions, thereby leaving information gated through task-relevant areas^69^. Consequently memory encoding and retrieval “benefit” from a desynchronized brain state reflected in reduction of alpha activity, as the storage capability of a system or “richness of information represented in the brain” was found positively correlated with the level of alpha power reduction^64, 72^. This conclusion, though, has the caveat of circularity: the cognitive construct, e.g. attention or WM, is associated with decrease in alpha power, which in turn is interpreted as reflecting the increase in attention or WM load. The oculomotor model overcomes this interpretational caveat. Directing attention to a single, couple of multiple items in an experimental task requires gaze control over (or away from) single, couple of multiple items. Consequently, the degree to which oculomotor action is controlled during these tasks manifests as fluctuations in cortical alpha activity.

Present results do not argue against definitions of the functional significance of alpha oscillations as formulated in the inhibition-timing^71^, gating by inhibition^69^, and desync hypotheses^72^. Central to these models is the role of alpha activity in the transition of external input (e.g. stimulus, event, memory) into internal cortical “representation”. Present results identify a lower level operation, prior to the manifestation of a higher-level cognitive construct, with high biological plausibility. It is the oculomotor action that allows this external-internal translation (i.e., focus on relevant - thereby neglect distracting - environmental items). Differential oculomotor action is involved in sensory recruitment in different working memory tasks (e.g. N-back vs. Sternberg) which resolves different alpha power modulation in these tasks: decrease with load in the former and increase with load in the latter (see Figure 6).

Spatial location is tracked by both alpha power modulation^8–10^ and alpha phase^73^. This conclusion from human studies parallels findings of place cells in the rodent literature^43^. Place cells are conceived as reflecting a fundamental mechanism translating to navigation not only in physical but in cognitive space (e.g., memory)^74^. Importantly, it is a mechanism that does not depend on external cues but occurs spontaneously^44, 45^. In contrast to the rodent literature, human findings on spatial attention have thus far been explained exclusively by the presence of a stimulus (cue) at a particular position. The reported evidence clearly indicates spontaneously generated patterns of oculomotor action that track position in physical space (see Figure 5). In line with observations made during visual and auditory spatial attention, spontaneously generated place topographies reveal the tiling of space with respect to the observing individual, i.e. an egocentric reference frame.

### Alpha phase determines the “Where” and “When” of oculomotor action

Human and nonhuman primate literature indicate involvement of alpha oscillatory phase in guiding saccadic behavior^37–39^. Alpha phase modulates saccade response latency^38, 40^, improves memory encoding by establishing connectivity between visual and memory circuits^37^, and maintains stable representation of the external world across eye movements^39^. With cortico-centric perspective (reviewed in ^75^), alpha phase is explained as a mechanism regulating neuronal excitability and connectivity between cortical areas during cognitive processing. Yet eye movements are somewhat neglected. Here, we presented evidence that oculomotor output beyond saccade onset (e.g., fixational tremor) is regulated by ongoing alpha phase. We conclude that the phase of alpha activity serves a dual “temporal” purpose. First, it aids *where (and how long)* to look at a given target by ensuring steady gaze position (Figures 4 and 7a). As a consequence, any potential input outside foveal vision is “suppressed”, which would be in line with an interpretation in the context of “inhibitory control”. Second, it determines *when (and how fast)* to saccade to a next target providing the temporal window utilized by SC in saccade generation. Rapid saccadic response toward a potential target benefits from a steady state reflected in the tremor regulation (Figure 7 b-d).

### Alpha oscillations support the cortical control of saccadic eye movements

As outlined in the introduction, research has identified key structures involved in oculomotor control distributed across visual, parietal, and motor cortex as well as the frontal eye fields^18, 19^. The latter considered as a key structure in visuo-spatial cognition (for a comprehensive review, see ^76^). Electrical stimulation of visual and parietal cortex and the FEF elicits saccades with similar amplitude and direction^19^. The firing rate of FEF neurons is increased just prior to saccade^77^ onset, which is in line with the present observation of increased alpha phase coherence in the FEF prior to saccade onset. Using paired visual targets presented at different levels of temporal asynchronies, lesion studies have confirmed that the FEF are involved in controlling a sequence of eye movements and the selection of visual targets^78^. FEF ablation leads to compromised selection of a visual target, highlighting the timing of saccadic eye-movements as one of the key functions of FEF. In individuals with left FEF ablation, the stimuli need to be presented ∼100ms earlier on the affected side in order to equalize the brain’s choice (left or right target selection). That is, a temporal window approximating the length of an alpha cycle suffices to achieve the same performance as in an intact individual. This is in line with studies using target-selection tasks, and indicates that visual areas and the FEF are functionally linked through alpha/beta activity^79–82^. The proposed theory linking alpha oscillations to oculomotor control offers an explanation of these results: the temporal window provided by the alpha phase is common to both the eye muscles (via the SC vector code) and the FEF, connected via the posterior system including parietal and visual areas ^82^. Thus, the link between visual cortex and FEF primarily serves oculomotor control in the decision process where to look next.

The involvement of the OFC prior to saccade onset is a finding that wasn’t anticipated. According to the existing literature, both the frontal scalp topography and the source localization of the saccadic spike artifact in the vicinity of the orbitofrontal areas^35, 83^ appears a plausible explanation for this finding as an ocular artifact. However, the time course of the alpha phase consistency, e.g. some 500 ms before the onset of the saccadic spike, argues against OFC activity as mere artifact. Alternatively, the OFC has been considered a structure within the cerebral circuit involved in saccade generation with the purpose of maintaining goal-related information (e.g., look toward or look away from a target; for review see ^84^). It is noteworthy that studies demonstrating increase in alpha inter-trial phase coherence prior to either a visual stimulus detection or saccade onset ^38, 85–87^ report consistent prefrontal topography, which is in line with this notion of prefrontal executive control in saccade generation. Another possibility is suggested by consideration of the relationship between saccades and pupil size. Saccades are routinely associated with changes in pupil size^88–90^, in turn a signature of sympathetic arousal and sensory gain. Pupil size indeed varied with saccade onset exhibiting transient decreases in pupil size after the saccade (supplemental Figure S3). Fluctuation in pupil size is regulated by cholinergic neurons residing in the nucleus basalis of Meynert (NBM) located in the basal forebrain, a location closely overlapping with the OFC location reported here. This cholinergic signaling propagates throughout the cortex ^91^ and has been hypothesized to be a key neuromodulator in the generation of alpha oscillations^92^. Indeed, the atrophy of basal forebrain nuclei has been associated with attenuation of posterior alpha oscillations ^93^. The modulation of cortical oscillatory activity has recently been highlighted in a study demonstrating covariation in mutual information between visual alpha/beta oscillations and pupil size^94^. As changes in pupil size is preceded by noradrenergic and cholinergic activity^95, 96^, the involvement of cholinergic circuits in the basal forebrain prior to saccade appears likely. The increase in alpha phase coherence reported here might, at least in part, reflect the electrophysiological manifestation of pupil size regulation. Nonetheless, a conclusion about the putative relationship, and its functional significance, between ongoing alpha phase in orbitofrontal circuits and saccade generation would be premature and should be addressed in future work.

### General conclusion

The present series of observations provides diverse support for the proposal that alpha oscillations link oculomotor action to cognition. The conclusion on a causal relationship is not supported by the present data and requires further investigation. Nonetheless, present results are in line with the notion that the main objective of brain computation is to maintain and control the input to its sensors^97, 98^. Saccades induce a transient gain in the primate brain^97^, as such a prerequisite for visually-guided cognitive processing.

## Acknowledgments

This work was supported by the Velux Stiftung Project No. 1126 and by the Schweizerischer Nationalfonds zur Förderung der Wissenschaftlichen Forschung (SNF) Grant 100014_175875 awarded to NL, the James S. McDonnel Foundation Understanding Human Cognition Collaborative Award 220020448 awarded to OJ and by the Deutsche Forschungsgemeinschaft (grant RO805/14-1,2) awarded to BR. The authors thank Gyorgy Buzsaki for stimulating discussions on earlier versions of this manuscript and all participants volunteering in the experiments.

## Online Methods

### Experiment 1

#### Human participants

Twenty-three male and 17 female individuals participated in the study. Age ranged between 22 and 41 (M±SD, 26.9 ± 5). Participants were recruited at the local university, reporting no history of neurological or psychiatric disorders. Written informed consent was obtained from each individual prior to participation. The study was approved by the University of Konstanz ethics committee.

#### Stimulus material and procedure

During a saccade calibration task, participants were positioned in front of a computer screen of 1920 pixels width and 1080 pixels height at a distance from the eyes of 116 cm. Participants were asked to move their eyes to visually follow the repositioning of a fixation cross as fast as possible. After a central fixation of 3000 ms, the fixation cross moved to a target location for 300 ms and back to central position. Thereafter, participants were allowed to blink for 500 ms before the onset of the next trial. Locations were chosen as follows: either left at −5 and – 10° and right at 5° and 10° horizontal eccentricity or up at 4° and 8° and down at −4° and −8° vertical eccentricity. Within eight successive blocks, ten repetitions were required for every location resulting in 80 trials in total. Stimulus presentation was controlled using Presentation software (www.neurobs.com) on a Windows 7 PC.

#### EEG acquisition and processing

EEG was recorded in an electrically shielded room with a 128 channel ANT Neuro system using equidistant hexagonal waveguard caps. EEG was sampled at 1024 Hz. Each channel was referenced online to the average. Electrode impedances were kept below 20 k∧. Continuously recorded data were epoched from 4000 ms before to 4000 ms after left or right saccade onsets. Subsequently, the exact sample points of saccade onsets were identified by visual inspection, and the data were re-segmented to adjust for the delay between the relocation of the fixation cross and the onset of a saccade, such that the 0 ms corresponded to saccade onset. The epoched data were then down-sampled to 300 Hz, and ICA was used to remove components associated with cardiac and eye movements. Data corresponding to left and rightward saccades (i.e., 40 trials) were considered for further analysis. Analysis was conducted using the open-source MATLAB toolbox FieldTrip^99^. Time frequency resolved power was computed for −2000 ms to 3500 ms (wider than the epoch subsequently analyzed, as required by the Hanning taper) and frequencies from 2 to 40 Hz in 2 Hz steps by convolving the data with a single 500 ms Hanning taper. Power estimates were expressed as change from pre-saccade baseline (−2000 to −500 ms) in units of decibel.

#### Primate data analyses

An openly available nonhuman primate data set was obtained from the neurotycho data base (http://www.neurotycho.org/fixation-task). During ECoG recordings of 128 implantented subdural electrodes, the participant was positioned in a primate chair with head and arms fixed. The participant was required to hold fixation on each trial with a fixation point presented at the center of visual display. At the timing of fixation disappearance, a reward was presented to the animal. Continuous EEG was sampled at 1 kHz and the corresponding eye-tracking data at 500 Hz. The recording was performed across several sessions on several days. The available data were already merged across sessions and epoched around the fixation point onset. In a first step, the ECoG data were downsampled to 500 Hz. Using the eye-tracking data, onsets of saccades were identified via visual inspection. The sample points of saccade onsets were then used as triggers to re-segment the data from 2000 ms before to 2000 ms after onsets. Subsequently, time-frequency analysis of power was conducted from - 1000 ms to 2000 ms following the procedures described above.

### Experiment 2

Three human pilot participants performed self-initiated leftward and rightward saccades with eyes closed. MEG was recorded at a sample rate of 1000 Hz using a 306-sensor TRIUX MEGIN system (MEGIN, Finland) in a magnetically shielded room (hardware filtering: 0.1-330 Hz). A signal-space-separation algorithm implemented in the Maxfilter program provided by the manufacturer was used to remove external noise (e.g., 16.6 Hz train power supply and 50 Hz power-line noise). Oculomotor events such as blinks and saccades were recorded using conventional vertical and horizontal EOG. EOG electrode impedance was kept below 10 kΩ. Saccade onsets were identified by visual inspection from the continuously recorded horizontal EOG and used as triggers to segment the data ± 4000 ms around saccade onset. Subsequently, the epoched data were down-sampled to 300 Hz. Time-frequency analysis was computed for −2000 ms to 3500 ms (wider than the epoch subsequently analyzed, as required by the Hanning taper) and frequencies from 4 to 40 Hz in 2 Hz steps by convolving the data with a single 500 ms Hanning taper. Time-frequency representation of power relative to pre-saccade baseline (−2000 to −500 ms) was expressed in units of decibel for occipital sensors (‘MEG2232+2233’, ‘MEG2312+2313’) as illustrated in Figure 2a.

### Experiment 3

#### Human participants

Twenty-four female and five male individuals participated in the study. Age ranged between 19 and 37 (M±SD, 21.8 ± 3). Participants were recruited at the local university, reporting history of neurological or psychiatric disorders. Written informed consent was obtained from each individual prior to participation. The study was approved by the University of Konstanz ethics committee.

#### Stimulus material and procedure

In a Posner prosaccade task, participants were instructed to maintain central fixation for 2000 ms. A cue was presented for 20 ms at one of seven possible locations (see Figure 2a and Figure S1) following by a 2000 ms delay interval. During this delay interval, participants were asked to maintain the location of the previously presented cue in memory, yet maintaining central fixation. Thereafter participants performed a saccade from central fixation toward the position of the cue. Each individual performed the experiment twice: once in a normal ambient light condition and once during full darkness. Order of participant assignment to light and dark condition was pseudorandomized. To ensure full darkness, new hardware was designed. Eight LEDs were soldered to an aluminum grid (390 mm width and 305 mm height) with positions illustrated in Figure S1. This hardware ensured full darkness in the Faraday cage during the off-periods of the LEDs, which is not the case with conventional computer screens, which emit some light when the screen is set to black. The central bottom position was left out because only eight positions were available to communicate with the trigger box. During EEG acquisition, participants were seated in an electrically shielded room with an approximate distance to central fixation of 50 cm to the eyes.

#### Data acquisition and preprocessing

EEG was measured in an electrically shielded room using a 256-channel HydroCel net (Electrical Geodesics, Inc., Eugene, Oregon, USA). EEG filtered 0.1 to 400 Hz was sampled at 1000 Hz. The Cz recording reference was changed to average reference in offline analyses. Electrode impedances were kept below 30 KΩ. Forty trials per LED position were acquired resulting in 120 left (up, center, down) and 120 right cued location. These 240 trials were kept for subsequent analyses. Data was segmented to include −2000 s to 4000 ms around cue onset. Following 45 Hz low-pass filtering and down-sampling to 200 Hz, ICA was employed for artifact control of saccades, eye blinks, and cardiac activity. No trials were removed. Electrodes nearest the neck and cheeks were excluded from further analyses to avoid contamination by muscular activity. Time-frequency-resolved power was computed for - 750 ms to 2000 ms (wider than the epoch subsequently analyzed, as required by the Hanning taper) and frequencies from 2 to 40 Hz in 2 Hz steps by convolving the data with a single 500-ms Hanning taper. Lateralization of alpha power estimates was expressed as effect size (Cohen’s d) based on the statistical contrast left vs. right (statistical analysis below).

### Experiment 4

#### Humanarticipants

A sample of 143 individuals (83 female) participated in the study. Age ranged between 19 and 83 (M±SD, 47.7 ± 22.7). Participants were recruited at the local university, reporting no history of neurological or psychiatric disorders. Written informed consent was obtained from each individual prior to participation. The study was approved by the University of Zurich ethics committee.

#### Stimulus material and procedure

In a prosaccade task, participants were asked to maintain central fixation for a jittered interval of 1000 to 3500 ms. Subsequently, a 1000 ms cue (i.e., dot) appeared horizontally either to the left or to the right (8° visual angle) of a central fixation square. Participants were asked to direct and maintain their gaze on the cue as quickly as possible. Prosaccade and antisaccade blocks were acquired. For the purpose of the present study only prosaccade data were analyzed. Two prosaccade blocks yielded a total of 120 trials (30 per hemifield) per recording session. Participants were examined during two recording sessions separated by a week in order to assess test-retest reliability.

#### Data acquisition and preprocessing

EEG was measured in an electrically shielded room using a 128-channel Geodesic Sensor NET (Electrical Geodesics, Inc., Eugene, Oregon, USA). EEG filtered 0.1 to 400 Hz was sampled at 500 Hz. The Cz recording reference was changed to average reference in offline analyses. Electrode impedances were kept below 40 KΩ. Electrodes nearest the neck and cheeks were excluded from further analyses to avoid contamination by muscular activity. Eye-tracking was recorded using the left eye with an infrared video-based eye-tracker (Eye Link 1000 Plus, SR Research, http://www.sr-research.com) with a sampling rate of 500 Hz. The calibration was done by a 9-point grid before every block of the prosaccade task. During the calibration, the participants were asked to direct their gaze toward nine targets appearing on the borders, the edges, and the middle of the screen. After the calibration, a validation was performed with the same protocol. The average error of all points was kept below 1°. The data used in this experiment were part of a larger project addressing test-retest reliability of EEG-derived metrics across different cognitive tasks. For reasons of reproducibility, a set of preprocessing steps common to all tasks was already in place and adopted in the current examination. In particular, EEG preprocessing was conducted with the MATLAB toolbox Automagic version 2.4.3^100^ and the MATLAB toolbox EEGLAB version 14.1.2b^101^. Bad electrodes were detected with the EEGLAB plugin clean_rawdata. Prior to bad electrode identification a temporary high-pass filter with a transition band of [0.25 0.75] Hz was applied with a line noise criterion set to 4 standard deviations. Furthermore, a correlation between the signal of each electrode and an estimate based on neighboring electrodes was calculated. If this correlation was below 0.85, the electrode was marked as bad. Residual bad electrode detection was done with a high variance criterion. Time points with an absolute amplitude higher than 100 μV were removed. If this process rejected more than 20% of the data, the electrode was marked as bad. For the remaining electrodes the standard deviation was calculated, and an electrode was removed if it exceeded 25μV. Zapline^102^ was applied to remove line noise artifacts, discarding 7 power line components. Additionally, an ICA with a temporary high-pass filter of 2 Hz was performed with the pre-trained classifier ICLabel^103^ to reduce non-brain artifacts. Each component classified with a probability of 0.8 for the artifacts of muscle activity and eye activity, heart artifacts, line noise, and electrode noise was removed from the data. Eventually, bad electrodes were replaced via spherically interpolation using the eeg_interp() function and rated with the objective quality criteria of Automagic. Any data file rated as bad, meaning that the proportion of high-amplitude data points in the signal (> 30 μV) was larger than 0.2, more than 20% of the time points showed variance larger than 15 μV across electrodes, 40% of the electrodes showed high variance (15 μV), or the proportion of bad electrodes was higher than 0.4 was not included in the analysis. After all the preceding artifact-correction steps, twenty-four electrodes nearest the neck and the face of the subjects were removed (E1, E8, E14, E17, E21, E25, E32, E48, E49, E56, E63, E68, E73, E81, E88, E94, E99, E107, E113, E119, E125, E126, E127, E128) to avoid contamination by muscular activity. Subsequently, average re-referencing was applied. The data were epoched from 2000 ms prior to 3000 ms after saccade onset. Epochs with incorrect responses toward the cue were rejected, as well as micro-saccades defined as saccades with an amplitude lower than 1.5° and response time lower than 100 ms. The information about the saccade for the error removal was derived from the eye-tracking system. Before EEG preprocessing, eye-tracking and EEG datasets were synchronized with the EYE-EEG toolbox^104^. Time frequency resolved power was computed for −2000 ms to 2000 ms and frequencies from 1 to 40 Hz in 1 Hz steps by convolving the data with a single 1000 ms Hanning taper. Power estimates were expressed as change from presaccade baseline (−800 ms to −500 ms) in units of decibel. The Hemisphere (L,R) × Saccade direction (L,R) time-frequency representation of power was computed as follows. First, alpha power lateralization was expressed as the difference between left and right saccades. Subsequently, three occipital electrodes over the right (‘E84’, ‘E85’, ‘E91’) and the left (‘E59’, ‘E65’, ‘E66’) hemisphere were averaged and subtracted yielding the time-frequency representation illustrated in Figure 3a and b.

#### Inter-trial phase coherence analysis

Time-frequency analysis was performed for the interval −2000 ms to 2000 ms around saccade onsets, and the Fourier coefficients were retained in the output. After normalization of each trial with its corresponding amplitude, the absolute value was computed and averaged across trials, reflecting the magnitude of inter-trial coherence (ITC). Reliability of ITC values form the test and retest recording sessions were evaluated.

#### Test-retest reliability estimates

Intraclass correlation (ICC) was computed using the agreement measure^105^. Both power and inter-trial phase coherence were evaluated using the data from combining the two recording sessions.

### Experiment 5

#### Participants

A publicly available MEG dataset of 89 human volunteers acquired during resting awake conditions was used. These data as well as all details about the participant sample can be found at https://www.humanconnectome.org/.

#### Data acquisition and preprocessing

Data were recorded using a whole-head, 248-channel magnetometer system (MAGNES 3600 WH, 4D Neuroimaging, San Diego, CA) with participants in the supine position. Data were continuously recorded with a sampling rate of 2034.5101 Hz and a bandwidth of DC-400 Hz. Digitization of the each participant’s head shape and of the locations of the fiducial coils was accomplished with a Polhemus 3Space Fasttrak system. Just prior to the working memory task (N-back paradigm), data from which were used in Experiment 6, the participant underwent three runs of approximately 6 min of eyes-open resting-state MEG recording, used in Experiment 5. The data from the three runs were appended and segmented into 1000 ms epochs. No further preprocessing was applied.

#### Forward encoding modeling

Forward encoding modeling followed the procedure described in^106^ (https://osf.io/vw4uc/). The general assumption is that alpha power quantified at each sensor reflects the weighted sum of eight hypothetical saccade directions reflecting the neural signal evoked by an eye movement in accord with the primary field of action of the ocular muscles, discussed in the main paper. Each of these hypothetical neuronal activation patterns is tuned to a different position/muscle direction. One-second epochs of MEG data were partitioned into two blocks (train and test) with equal epoch numbers. A ten-fold random generation of multiple block assignments (e.g., test or train) was utilized, and the outcome was averaged over folds. Single-epoch alpha power was estimated using a Fast Fourier Transformation (FFT) after multiplication with a single Hanning taper. A set of eight basis functions coding for eight equally spaced positions (P1 to P8 in Figure 5) between 0 and 360° was constructed. Training data *B1* allowed the estimation of weights that approximated the relative contribution of the eight hypothesized tuning functions (*k*) to the measured scalp data. Let *B_1_* (*m* sensors × *n_1_* epochs) be the signal at each electrode and epoch in the training set, *C_1_* (*k* tuning functions × *n_1_* epochs) the predicted response of each tuning function, and *W* (*m* sensors × *k* tuning functions) the weight matrix allowing the linear mapping from the tuning model to MEG sensor space. This mapping was based on a linear model of the form:

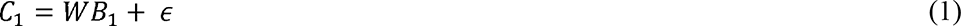

Where ∈ contains (assumed Gaussian) error terms that should be minimized. Ordinary least-squares regression was used to estimate the weight matrix *W (m × k)*:

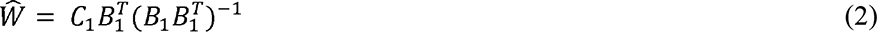

On the basis of this weight matrix and the test data *B_2_* (*m* sensors × *n_2_* trials), an estimated response matrix *C_2_* (*k* tuning functions × *n_2_* trials) was calculated:

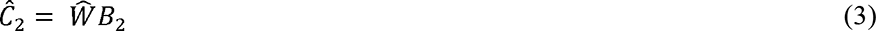

The estimated responses were circularly shifted by degrees such that estimates associated with positions that evoked a response were centered at 0° of the position space (P1 to P8) spanning −180° to 180°. Following this step, an accurate model is characterized by a maximum at 0° and a minimum at −180°/180°. In contrast, an inaccurate model fit approximates a flat line. This was repeated until each block had served as a training set and a test set.

Finally, to interpret the weight matrix *W* in terms of source origin, an activation pattern represented in matrix *A* of a corresponding forward encoding model was computed^107^:

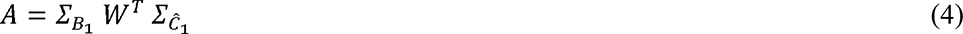

Here, 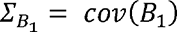 and 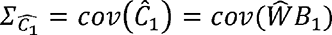 are covariance matrices. The advantage of using *A* instead of the raw weights *W* is that elements of *W* may reflect suppression of “signals of no interest”^107^. For example, correlations across sensors in *B_1_* could be confounded by noise. Therefore, they do not reflect brain activity related to *C_1_*. Transforming to activation patterns *A* mitigates this problem. This forward encoding model was applied 1000 times for low vs. high alpha trials separately, determined by median split for each subject. At each iteration, each trial was randomly assigned a hypothetical position between P1 and P8. The results presented herein were averaged over the 1000 iterations.

### Experiment 6

Results presented in Figure 5 are based on publicly available datasets. These include: a) data from a study examining working memory (WM) in the context of the Sternberg task published in^57^ and accessible at https://osf.io/w6d92/, b) data provided by the human connectome consortium including participants performing the N-back WM task published in^54^ and accessible at https://www.humanconnectome.org/, c) same resource as in b) but for the resting-state eyes-open data, d) data provided by www.neurotycho.org with details about the animal recordings and task description listed at http://wiki.neurotycho.org/Fixation_Details, and e) data from a study examining odor-induced oscillatory activity in honey bees^59^ and accessible at https://osf.io/523tk/?view_only=bfa999c448014bd196d1f432fb3c593e. Details of participants and task procedures are available in the aforementioned papers.

#### Data preprocessing and analysis: Figure 6a

Raw data from participant S06 reported in^57^ was evaluated. This participant was chosen based on the clear modulation of alpha power with memory load as frequently reported in the literature. The data were epoched from −2000 to 3000 ms following the onset of the maintenance interval of WM items. ICA decomposition was used to identify and reject activity associated with eye blinks after which the data were re-segmented to include the maintenance interval of 3000 ms. A spectral decomposition using FFT and multitapers was applied for the frequency interval of 4 to 40 Hz and frequency resolution of 0.33 Hz. EOG data were extracted from two electrodes position in the vicinity of the left and right canthi respectively (‘LD2’ and ‘RD2’) and re-referenced as a bipolar channel. These horizontal EOG data were spectrally decomposed in the frequency range of 0 to 200 Hz, as it is well documented that muscular activity covers a broad range of frequency. Finally, behavioral results (reaction time), alpha power, and oculomotor activity were illustrated (Figure 6a).

#### Data preprocessing and analysis: Figure 6b

Raw data from 83 participants performing the N-back task was used (37 female, mean age 28.5 years, range 22-35). Data were segmented from 0 to 2000 ms following the target stimulus. No further preprocessing was applied. Spectral decomposition of the MEG data was performed from 3 to 40 Hz and for the horizontal EOG from 0 to 300 Hz. Grand-average waveforms and results are provided in Figure 6b.

#### Data preprocessing and analysis: Figure 6c

Procedures identical to those for Figure 6b were applied to the resting-state data from 89 participants. MEG and horizontal EOG data were read in continuously and re-segmented into windows of 1000 ms. Spectral decomposition was applied (3 to 40 Hz), and indices of trials with high vs. low alpha power were computed based on median split for each participant. Grand-average waveforms and results are provided in Figure 6c.

#### Data preprocessing and analysis: Figure 6d

The primate data as preprocessed for Figure 1 were used. First, alpha power (9-14 Hz) was estimated using FFT analysis for two electrodes located in the vicinity of area LIP (electrodes ‘45’ and ‘46’). Indices of trials dominated by high vs. low alpha power were computed based on median split for each subject. Spectral analysis of the entire data was performed in the range 3 to 40 Hz and subsequently segmented into low and high alpha power conditions. For the same trial indices, eye tracking data were spectrally analyzed in the frequency range 0 to 200 Hz. Finally, cortical alpha power and power derived from the horizontal eye movement data were illustrated by condition (high vs. low alpha power). The data from five honey bees (Apis mellifera) were used. Time-frequency analysis from - 3750 to 3750 ms around odor onset was performed using a sliding window of 500 ms covering the frequency range from 2-40 Hz in steps of 2 Hz. The animals were examined while the sensory antenna was amputated on the side resulting in an ipsilateral reduction in ongoing oscillatory activity (Figure S1 in ^59^). This effect is illustrated in the Figure 6d.

### Experiment 4 - procedures related to Figure 7

A single participant from the prosaccade task sample described above was evaluated in the initial exploratory analysis illustrated in Figure 6. The continuous data weresubjected to an ICA decomposition. Components associated with posterior alpha activity were selected by visual inspection based on 1) clear posterior topography and 2) time course dominated by alpha activity. The resulting EEG data were bandpass filtered 7-14 Hz and plotted together with the time course of the horizontal eye-tracking data (Figure 6a). These eye-tracking data were high-pass filtered at 40 Hz for illustration of the alpha activity and eye tracking illustrated in Figures 6b and 6d. Cross-frequency coupling analysis between the phase of ongoing EEG alpha activity and the amplitude of the high-frequency activity in the horizontal eye-tracking data was performed using the modulation index method^108^. Time onsets of saccades were identified and replaced with zeros using the *ft_artifact_zvalue* function in FieldTrip. This step was necessary to account for the large “jumps” associated with large saccades, in turn artificially contributing high energy in the frequency spectrum. For phase in frequencies in the range between 2 to 16 Hz in steps of 2 Hz, time-frequency decomposition was performed using a single Hanning taper and a sliding window of 2 Hz. High amplitude providing frequencies were estimated using multitapers^109^ in the frequency range 30 to 120 Hz. Finally, this exploratory cross-frequency analysis was extended to an additional 137 human volunteers, and the test-retest reliability was computed.

### Statistical analyses

Inferential statistics were carried out using a cluster-based approach based on randomization^110^. This approach identifies clusters (in time, frequency, and space, i.e., electrodes) of activity on the basis of which the null hypothesis can be rejected, while addressing the multiple-comparison problem. A two-tailed alpha threshold of 0.05 and 1000 permutations were used. t-values were converted to Cohen’s d effect size.

### Source-space analyses

Source estimates were computed in the frequency (Experiment 1) as well as in the time (Experiment 4) domains. In the frequency domain, an adaptive spatial filtering algorithm was used (dynamic imaging of coherent sources, DICS)^111^. This algorithm uses the cross-spectral density matrix from the EEG data and the lead field derived from the forward model to construct a spatial filter for a specific location. This matrix was calculated using a multi-taper FFT approach for data 200 to 1200 ms following saccade onset. Spectral smoothing of ±2Hz around center frequencies of 10 Hz was applied including the power in the 8-12 Hz (alpha) range. These spectral density matrices and thus the spatial filters were participant-specific and estimated based on pre-saccade-onset and post-saccade-onset data. This so-called common spatial filter ensures that potential differences in oscillatory power between pre-saccade and post-saccade intervals are not due to differences in filter estimates (e.g., resulting from different SNR in pre-saccade vs. post-saccade).

In Experiment 4, the linearly constrained minimum variance beamforming algorithm (LCMV)^112^ was used. When applied to the data covariance matrix and forward models of the locations of interest, these algorithms construct spatial filters that allow the transformation of scalp-level data to source estimates at a given brain location of interest. Forward models were derived from the MNI ICBM 2009 template brain (http://www.bic.mni.mcgill.ca/ServicesAtlases/ICBM152NLin2009) using the OpenMEEG implementation^113^ of the Boundary Element Method (BEM). A parcellation scheme based on the Desikan-Kiliani atlas^114^ was used. This parcellation scheme consist of 34 cortical ROIs per hemisphere. Following procedures similar to those described in ^115^, single-dipole-specific spatial filters were concatenated across vertices comprising a parcel, yielding a set of 34 time courses of activity per hemisphere. For each parcel, a principal component analysis was performed to extract spatially orthogonal and temporally uncorrelated components that describe the activity, ordered by the amount of variance explained. The first principal component was selected as the representation of the parcel’s time course of activity. Finally, these source time courses were used to derive ITC on the source level as described above.

## Data availability

Analysis code to reproduce all reported results as well as the data that has not been already publicly shared will be made available at https://osf.io/.

