## Supplemental Figures for "Alpha oscillations link action to cognition: An oculomotor account of the brain’s dominant rhythm"

### Supplemental material

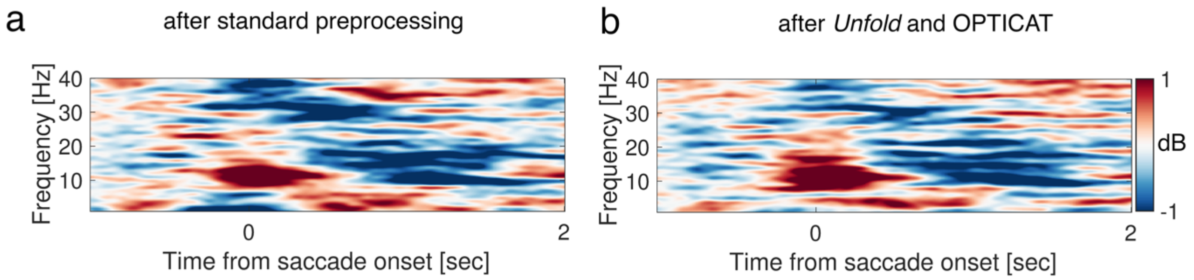


***Figure S 1: a-*** *Time-frequency representation of power (TFR) as in Figure 3 grand averaged over a subsample of 14 participants.* ***b****- TFR of the same participants after correcting the time series for ocular artifacts using the EEG deconvolution toolbox[1] Unfold and applying optimized ICA training (OPTICAT)[2] to remove residual myogenic artifacts.*

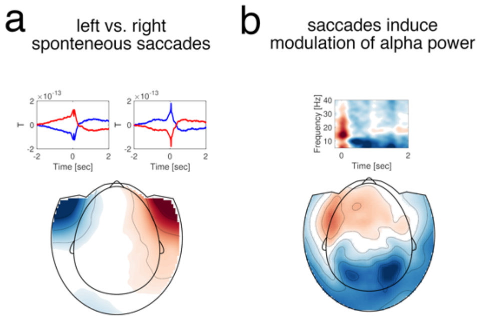


***Figure S 2: a-*** *Control analysis ensuring that saccades directed toward the left or right visual field are present in the spontaneous recordings. The topography represents MEEG for left minus right saccades around saccade onset at 0 sec. Time courses in the insets represent the activity over left frontal sensors (left, and blue region in the topography) and right frontal sensors (right, and red region in the topography). Blue line denotes leftward and red line rightward saccades. Note that saccade size exhibits considerable variation thus is not clearly reflected in the average. The saccadic spike response at 0 ms is however preserved.* ***b-*** *Control analysis replicating that alpha power is smaller after spontaneous saccades relative to pre-saccade baseline with clear occipital-posterior topography.*

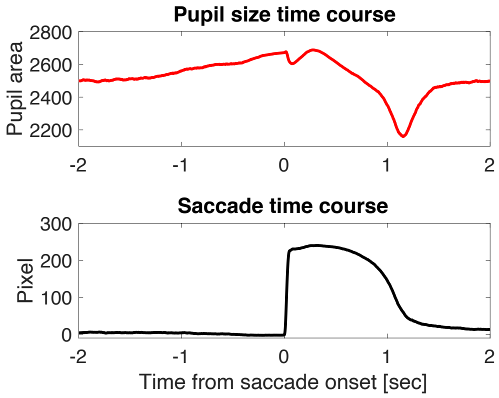


***Figure S 3: Top-*** *grand averaged time course of pupil size change relative to saccade onset at 0 sec.* ***Bottom****- corresponding grand average of saccade time course.*

1 Ehinger, B. V. & Dimigen, O. Unfold: an integrated toolbox for overlap correction, non-linear modeling, and regression-based EEG analysis. *PeerJ* **7**, e7838, doi:10.7717/peerj.7838 (2019).
2 Dimigen, O. Optimizing the ICA-based removal of ocular EEG artifacts from free viewing experiments. *Neuroimage* **207**, 116117, doi:10.1016/j.neuroimage.2019.116117 (2020).
